# Aberrant neuronal differentiation and splicing defects in Congenital Myotonic Dystrophy (DM1) iPSC models

**DOI:** 10.64898/2026.06.25.734569

**Authors:** Surya Chandra Rao Thumu, Jean Patrick Gonzales, Soha Munir, Connor Tuck, Oscar Dominguez, Sandeep K. Singh

**Author notes:** Correspondence: Dr. Sandeep K. Singh, PhD, Email.

## Abstract

Myotonic Dystrophy type 1 (DM1) is an autosomal multisystem disorder manifested due to unstable CTG nucleotide repeat expansion within the 3′-untranslated region of the dystrophia myotonica protein kinase (*DMPK*) gene. Although progress towards understanding of molecular pathogenesis in muscle and heart has been made, the pathways that affect the brain in DM1 is fundamentally unknown. In addition, the congenital DM1 manifest even more complicated brain abnormalities. Despite the wealth of existing cellular and animal models, iPSCs based studies are being fostered as they replicate the human model more closely to the disease. In view of this context, we set out to characterize the differentiation potential of congenital DM1 patient derived iPSC lines towards neuronal cells. Using neurogenin2 (NGN2) induced direct reprogramming of iPSCs into neurons and chemically defined media-induced neural induction protocol, we find that congenital DM1 mutant iPSC derived neurons exhibited precocious differentiation, as evidenced by their expression of pan-neuronal markers TUJ1 and Map2, along with increased processes extension and neurite length. Moreover, unbiased RNA sequencing analyses and qPCR validation revealed precocious and enhanced expression of several neurogenic transcription factors including, Ascl1, NeuroG2, and NeuroD1. Furthermore, immunofluorescence imaging of MBNL1 and MBNL2, RNA-splicing factors, displayed enhanced nuclear aggregations, a hallmark of the DM1 disease, in the mutant lines. Moreover, investigation of RNA splicing events identified mis-splicing in many important genes/transcripts including RMST, ANK3 and MBD1 during the neural conversion of congenital DM1 lines. These studies reveal novel paradigms that may contribute to neurological pathogenesis in CDM1 patients. These studies also provide a strong foundation for future mechanistic investigation aimed at understanding CDM1 pathology and may open new avenues for the development of gene therapy approaches for individuals with DM1.

## Introduction

Myotonic dystrophy (DM) is a neuromuscular disease characterized by autosomal dominant inheritance with the global pooled prevalence estimated to be 9.99 per 100,000 (Liao et al. 2022). It affects primarily the skeletal muscles leading to myotonia, progressive muscle weakness and wasting. In addition, it affects multiple organs including the brain and is characterized with plethora of clinical symptoms involving early cataracts, cardiac conduction abnormalities, abnormal glucose tolerance, gastrointestinal dysfunction, cognition disabilities, excessive daytime sleepiness, frontal baldness, choking or coughing during swallowing and dyspnea (Meola and Sansone 2007; Wu et al. 2024). With the discovery of specific gene mutations, DM has been categorized as DM type 1 (DM1) and DM type 2 (DM2). DM1 also known as Steinert’s disease is caused by the presence of unstable CTG trinucleotide repeats (ranging from 37 to several thousands) in the 3’ untranslated region of myotonic dystrophy protein kinase (DMPK) gene located on the chromosome 19q13.3 (Brook et al. 1992). The severity of the symptoms is correlated with the number of CTG repeats in such that longer CTG repeats are associated with onset of the symptoms at an earlier age and with more rigor contributing further to its impressive variability (Echenne and Bassez 2013).

Although myotonic dystrophy has long been called the muscle condition, CNS related conditions are observed from birth to adult stages in DM1 patients. The cognitive disabilities increase with age and worsen in patient individuals with the increased number of CTG repeats. Clinically five distinct and overlapping forms of DM1 have been categorized based on the onset of the disease: the congenital form (from birth to one month), the infantile (between 1 month and 10 years old) the juvenile form (age between 10 and 20 years old) adult form (between 20 and 40 years old) and the late-onset form (onset after 40 years old) (De Serres-Berard et al. 2021). Among these, the congenital form (CDM1) is characterized by severe hypotonia, respiratory insufficiency and intellectual disabilities. Amidst the clinical manifestations related to CNS, the cognitive impairment is one of the important hallmarks of CDM1. It has been characterized with diminished capacity for visuospatial creation, executive function impairments like attention deficit disorder, and emotional or mental conditions including anxiety, mood disorders, phobias (reviewed in (Douniol et al. 2012; De Serres-Berard et al. 2021) and autism spectrum disorder and its traits (Angeard et al. 2018) (Sznajder et al. 2025) (Ekstrom et al. 2008; Patel et al. 2024). Histological analysis revealed ventricular dilatation, periventricular white matter lesions, and neuronal migration defects such as neuronal heterotopia and the presence of mature neurons dispersed randomly in white matter (Garcia-Alix et al. 1991; Endo et al. 2000; Meola and Sansone 2007; De Serres-Berard et al. 2021) (Hageman et al. 1993). Notably the manifestation of ventriculomegaly in the prenatal stages approximately corelates with the cognitive defects in young CDM1 patients (Martinello et al. 1999; Mutchnick et al. 2016; De Serres-Berard et al. 2021). While the presence of inclusion bodies and neurofibrillary tangles (NFTs) are confined to older DM1 patient brains (Hageman et al. 1993; Oyamada et al. 2006; Itoh et al. 2010); it suggests adult DM1 is a neurodegenerative condition while congenital DM1 is a developmental disorder. Despite important discoveries such as the involvement of CTG repeats in DMPK gene, dysfunction of MBNL family and other RNA binding proteins and dysregulated alternative splicing events in DM1(Wang et al. 2007; Batra et al. 2014; Goodwin et al. 2015; Tahraoui-Bories et al. 2023), the infantile onset of brain abnormalities in CDM1 remain uncertain. This raises the question of potentiality of therapies in development for treating this form of the disease.

To understand the effects of CTG repeats of DM1 *in vivo*, several mouse models were generated. Of these, the DMSXL mouse model was created with more than 1000 CTG repeats of the DMPK gene that manifest abundant nuclear foci and severe symptoms including retarded growth and high perinatal mortality representing early-onset DM1(Huguet et al. 2012). However, DMSXL model or MBNL1/2 double knock-out mice which were used for studying the neurodevelopmental disorders, could not recapitulate the CDM1 clinical manifestations such as ventricular dilatation (De Serres-Berard et al. 2021). This indicates a limitation to encapsulate the exact developmental disorders in humans thus confronting the advancement of treatment strategies. One of the powerful tools to study the molecular mechanisms of human CNS disorders is to use human induced pluripotent stem cells (iPSCs) and differentiating them to neurons *in vitro* (Ochalek et al. 2017; Sison et al. 2018; Kalia et al. 2023; Kern et al. 2025). The potential of iPSCs for their self-renewal and capability to differentiate into any adult cell in contrast to embryonic cells offers an advantage in CDM1 studies. The advances in iPSC biology have powered the possibility of generating cell types that are even impossible to access in patients. This has promoted for perusing the molecular mechanisms that are distinct in individual patients and offer personalized therapy. In this study, we exploited the stem cell technology to generate neurons from iPSC lines derived from CDM1 patients and a control individual. We employed the lentivirus mediated NGN2-induced direct reprograming and conventional chemically-defined medium-based approaches in differentiating the iPSCs to neurons and found mutant CDM1 lines to possess extensive neuronal differentiation behavior. We also unveil the dysregulated expression of several bHLH and other transcription factors (TFs) that favor neuronal switch. In addition, we found that the premature neuronal formation is accompanied with nuclear aggregation of RNA binding proteins MBNL1 and MBNL2 and several deficits in alternative splicing events.

## Materials and methods

### Maintenance and culturing of iPSCs lines

The iPSC lines of control (CTL) and the CDM1 (CDM1A and CDM1B) were obtained from **Neurology department, VCU**. iPSCs were maintained as feeder free cells in mTeSR medium (Stem cell Technologies) in 6 well plates (Thermo) coated with Matrigel (Corning) in 1X DMEM/F12 medium according to manufacturer instructions. During the cell maintenance, the growth medium is replaced with fresh medium daily. For storing the cells in liquid N_2,_ the cells are allowed to reach 70% confluency, rinsed gently with warm 1X DPBS -MgCl_2_, -CaCl_2_ (Thermofisher) followed by treatment with warm Accutase (Millipore) for 5-7 min. When the cells are observed detaching from the wells, accutase is diluted with 1X DPBS (-MgCl_2_, -CaCl_2_) and gently transferred to centrifuge tubes without trituration to maintain the iPSCs in clusters. The cells are centrifuged at 300 X g for 5 mins. The supernatant is removed, and the cell pellet is resuspended in freezing medium (mTeSR medium containing 10% DMSO) by gentle trituration only once or twice and aliquoted in 2 X 10^6^ cells per ml in freezing vial and stored in -80 °C and transferred to liquid N_2_ the following day.

### Southern Hybridization

For the detection of CTG repeats from the iPSC and control lines, 5ug of the isolated genomic DNA was digested with HaeIII and AluI restriction enzymes (New England BioLabs) and separated on 7.5% agarose gel. The gel was soaked in 0.25N HCl solution for 10 mins with gentle shaking followed by 2x denaturation buffer (0.5M NaOH, 1.5M NaCl). The gel was blotted onto nylon membrane in Southern apparatus using upward capillary transfer system overnight. The membrane with the transferred DNA is rinsed in 2x SSC buffer for 2mins on orbital shaker and allowed to dry at room temperature (RT) for 10 mins on 3mm Chr Whatman filter paper and baked at 120°C for 20 mins. Post baking, the membrane is rinsed in 2X SSC buffer for 1 min and suspended in buffer (5x SSC, 0.1% w/v N-laurosyl sarcosine, 0.02% SDS, 1% DMSO) for 1h at 70°C in rotator oven. The prehybridization solution is replaced with hybridization solution containing denatured DIG labeled locked nucleic acid (LNA) probe (5’ = /5DigN/GC+A G+CA GC+A G+CA G+CA GC+A GCA -3’) and incubated at 70°C for 4h. The membrane is washed 3x in low stringency buffer and followed by washing with maleic acid buffer containing 0.2% Tween 20. The membrane is blocked using blocking buffer (5 g Blocking Reagent (Roche, Cat. 11096176001) and Qs to 50 m with maleic acid buffer) for 30 mins and probed with DIG antibody.

### Lentivirus preparation

Lentiviruses were produced in HEK293T cells grown/maintained in DMEM containing 10% fetal bovine serum (FBS) by co-transfection with three plasmids: a packaging plasmid (psPAX2), envelope plasmid (VSV-G) and a vector plasmid (FUW-rtTA (addgene#20342) or FUW-TetO-Ng2-puromycin (addgene# 52047)). Packaging plasmid (15.75 μg), envelope plasmid (5.25 μg), and vector plasmid (7.00 μg) were co-transfected per one 150 mm culture dish using polyethyleneimine (PEI) of 1mg/ml stock. The PEI:DNA with a ratio of 3:1 is mixed and allowed to stand at room temperature (RT) for 10 mins and added to the cells and mixed gently. After 7-12 hours of transfection, the media containing the transfection mixture was removed and exchanged with prewarmed fresh complete medium and incubated further. The virus supernatant was collected every 24, 48 and 72h and stored at 4 °C. The collected supernatant at different timepoints was pooled, centrifuged for 10mins at 1000 rpm, filtered through 0.45 um filters and pelleted by ultracentrifugation at 23,000 rpm for 3h at 4 °C (**Beckman Coulter**). The viral pellet obtained was resuspended in ice cold 1X PBS, aliquoted and stored at −80 °C for future use.

### Isolation of astrocytes from mouse brain

Mouse cortical astrocytes were obtained from the newborn P3 wild type pups as described earlier with certain modifications (Gonzales et al. 2026). Briefly, the cortices from the pups were micro dissected and the tissue is digested with papain in the presence of DNase at 36° C for 45 mins with gentle swirling at every 15 minutes. This was followed by trituration with low and high ovomucoid solutions. The cells were passed through 20 µm mesh filter and resuspended in astrocyte growth media (DMEM suspended with 100 U/mL Pen/Strep, 2 mM L-Glutamine, 1 mM Na Pyruvate and 10% FBS) and 10-14 million cells were plated in 75 mm^2^ flasks coated with poly-D-lysine and incubated at 37°C in 10% CO_2_. All the media components were procured from Gibo unless otherwise stated. On DIV3, the growth medium was removed and rinsed with DPBS (-MgCl_2_ and -CaCl_2_) and the flasks were shaken by hand for 10-15 seconds or until a monolayer of astrocytes remained. The cells were rinsed with DPBS (-MgCl_2_ and -CaCl_2_) once and replaced with fresh growth medium. Once the astrocytes reached confluency, cells were trypsinized with 0.25% trypsin and passaged to 6 well dishes with 7-8 million cells per well.

### Induced neurons (iN) from iPSCs using Ngn2 transcription factor

Direct conversion of iPSCs to neurons was followed as previously described **(Zhang et al. 2013)**. Briefly, on day −2, iPSCs grown in mTeSR medium were rinsed once with DPBS (-MgCl2 and -CaCl2), treated with Accutase and plated as dissociated single cells onto Matrigel (Corning) coated 6 well (1×10^6^ cells/well) and 8 well chambered slides (4 X 10^4^). On day −1, lentivirus was suspended in fresh mTeSR^™^1 medium (1μl /ml) along with polybrene (8 μg /ml), mixed well and added to the cells for transduction. On day 0, the culture medium was replaced with N2 supplement/DMEM-F12 containing NEAA (1%), human BDNF (10 ng/ml, PeproTech), human NT-3 (10 ng/ml, PeproTech) and mouse laminin (0.2 ug/ml, Invitrogen). Doxycycline (10 μg/ml, Thermo-Scientific) was added to induce TetO gene expression and retained in the medium until the end of the experiment. On day 1, a 24 h puromycin selection (1 μg/ml) period was started. On day 2, mouse astrocytes isolated above were trypsinized, cell counts were taken and suspended in Neurobasal medium supplemented with B27/Glutamax (Invitrogen) containing BDNF, and NT3. Cell numbers of 5 x 10^4^ and 1.5 x 10^4^ were added to 6 well and 8 well chambered slides respectively. After day 2, 50% of the medium in each well was exchanged every 2 days. FBS (2.5%) was added to the culture medium on day 10 to support astrocyte viability, and iN cells were assayed on day 14 or 21 in most experiments.

### Neural induction of iPSCs to neurons

For neural induction of iPSCs, cells maintained to confluency in mTeSR medium were Accutase treated and triturated to single cells and centrifuged at 300Xg for 5 mins. The cells were resuspended in neural induction medium (NIM) containing SMAD inhibitor (STEMCELL Technologies) and only on the day of seeding, 10 mM Y-27632 (ROCK1 inhibitor) was added for the cells to attach. Cell counts were taken from the resuspended cells and seeded 2.5 x 10^6^ cells per well onto Matrigel precoated 6 well plates, 0.5 x 10^6^ cells per well of 12 well plates and 0.05 x 10^6^ onto 8 well chambered slides (for immunostaining). The NIM was replaced with fresh medium on daily basis. After 1 week (P#1), the cells in 6 well dishes were accutase treated, and replated onto fresh Matrigel coated plates for P#2 and the same was repeated until 3^rd^ week (P#3). At the end of every week, RNA was isolated from cells seeded in 12 well plates and the cells plated onto 8 well chambered slides were subjected to immunostaining.

### Immunostaining of differentiated neurons

For immunostaining, cells were fixed in 4% paraformaldehyde (PFA) prepared in sterile 1x PBS from 16% PFA solution (EM grade, Electron Microscopy Sciences) for 10 minutes. Following fixation, cells were rinsed with sterile 1x PBS twice at room temperature (RT) followed by blocking with blocking buffer containing 50% Natural Goat Serum (NGS) blocking and 0.2% Triton-X100. Following this, the cells were incubated overnight at 4°C with the following antibodies in blocking buffer containing 10% NGS: anti-MBNL1 (1:50) and anti-MBNL2 (1:50), anti-Map2 (1:2000,) anti-NeuN (1:1000,) anti-synapsin (1:1000,) anti Tuj1 (1:2000,) anti-Musashi (1:1000,) anti-Sox2 (1:500,). Subsequently, the cells were washed 3x with 1X PBS and incubated with species specific secondary antibodies conjugated with Alexa Flour 488 (1: 500) or Alexa Flour 594 (1:500) in blocking buffer for 1h at RT. This was followed by washing 3x with 1X PBS and finally mounting with DAPI containing vectashield mounting media (Vector labs). Cells were imaged as 20X and 63X images using confocal microscope (Zeiss LSM 710)

### Real-Time PCR

Total RNA was prepared from cells lysed using Trizol (Life Technologies). RNA was isolated from cells using Kit method (Machery-Nagel) as per the manufacturer instructions. The RNA was treated with DNAse and is reverse transcribed with a cDNA kit (Applied Biosystems) and amplified on the BioRadCFXConnect Real-time System. Species (human) specific pre-designed SYBR green primers (Bio-Rad) were used for PCR amplification. Gene expression levels were normalized to either TUBB3 mRNA and presented as fold expression over the control.

### RNA quality control, RNA-seq library preparation, sequencing and RNA-seq analyses

Total RNA isolated using Trizol method from above was used for RNA sequencing. RNA quality was assessed using a Fragment Analyzed 5200 system. All the samples showed RNA integrity number (RIN) of more than 7, indicating high RNA quality suitable for RNA sequencing. RNA-seq libraries were prepared from total RNA using the NEBNext rRNA depletion kit and NEBNext Ultra II directional RNA library prep kit for Illumina. Libraries were sequenced as paired-end reads on an Illumina NextSeq 2000 platform. Sequencing depth exceeded 40 million reads per sample.

Transcriptomic differences between samples were assessed using principal component analysis (PCA) based on normalized gene expression data. Correlation analyses were performed to evaluate relationships between control and CDM1 samples. Distinct clustering patterns confirmed separation of the three cell lines based on global gene expression profiles. For downstream analyses, RNA-seq datasets from passages P1, P2 and P3 were combined for each cell line to increase statistical power and enable robust comparisons across experimental groups, including aggregated induced pluripotent stem cell (iPSC) datasets.

### Alternative splicing analyses

Alternative splicing analysis was performed using the replicate Multivariate Analyses of Transcript Splicing (rMATS) framework to identify differential splicing events between experimental conditions. The analysis was based on a two-group comparison design, defined as Sample 1 and Sample 2 corresponding to control (CTL) and case (CDM1 or CDM2) respectively. rMATS was used to quantify exon exclusion levels and to identify differential alternative splicing events between groups. Splicing changes were evaluated using percent spliced-in (PSI) metric, and differences between CTL and CDM1 (averaged PSI of CDM1A and CDM1B)) were calculated as the inclusion level differences (ΔPSI) where CDM1 and CDM2 PSI were averaged for the final calculation. Statistical significance was determined using false discovery rate (FDR) to correct for multiple testing. Events were considered significantly differentially spliced if they met the thresholds of FDR< 0.05 and |ΔPSI| ≥ 0.1. To further improve the robustness and reduce artefacts associated with the low read coverage, an additional filtering step was applied. Splicing events were required to have a minimum of 5-10 inclusion and exclusion junction count in at least 70% of samples within each group. This criterion was used to exclude events with low read support that can yield artificially extreme PSI estimates.

### Gene ontology and PCR for alternative splicing (AS) analysis and

From the list of genes obtained from sequencing results of differential AS events, primers were designed between the upstream end and downstream start regions of the selected exons. The preferred list of genes and the primer sequences are provided in Table 1. The cDNA derived from the RNA extracted during the neural induction was used as the template. The PCR reaction was caried out at 94°C for 4 mins, 35 cycles of 94°C for 45 sec, annealing temperature (Table 1) for 45sec and 72°C for 30 sec. The amplified products were run on 2% agarose gel and captured. A total of 392 genes that fall into mis-spliced gene category between the control and mutant during RNA sequencing were subjected to gene ontology (GO) analysis. GO was performed using ShinyGO 0.85 version (https://bioinformatics.sdstate.edu/go/).

### Western Blotting

The cells were lysed in ice-cold modified RIPA buffer (20 mM Tris pH 7.4, 150 mM NaCl, 1 mM CaCl_2_, 1 mM MgCl_2_, 1% Triton-X100, 0.5% NP-40 and protease/phosphatase inhibitor) followed by sonication and centrifugation at 12,000 rpm for 15 min at 4°C. The supernatant was collected into a fresh tube and subjected to protein quantification using BCA protein assay kit (Pierce; #23227), and equal amounts were separated by SDS-PAGE on 10% gels and transferred onto nitrocellulose membranes (Bio-Rad). Blots were first blocked in 10% blotting grade milk and then incubated with the following primary antibodies: anti-MBNL1 (1:100) and anti-MBNL2 (1:100). Immuno-positive bands were visualized on automated machine by chemiluminescence with horseradish peroxide conjugated secondary antibodies (Jackson ImmunoResearch Laboratories, West Grove, PA, USA) and Pico Plus substrate (Thermo Fisher Scientific). Actin band intensity was used to normalize the expression levels.

### Quantification and statistical analysis

Sample sizes and statistical tests can be found in the accompanying figure legends. Offline analysis was performed using Graphpad Prism 11 and Microsoft Excel. Statistical significance was determined two-way ANOVA for comparison of 3 groups. Data are presented as mean□±□s.e.m. Statistical significance is indicated by asterisks: \**P□*<□0.05, \*\**P□*<□0.01, \*\*\**P□*<□0.001, \*\*\*\**P□*<□0.0001. The number of independent cultures (*n*) is noted in the figure legends.

Although majority of data is from three independent cultures, the differentiation was performed multiple times in due course of > 2 years of time with similar results of precocious neuronal differentiation by CDM1 mutant cells.

## Results

### Characterization of control and CDM-1 mutant iPSC lines

iPSCs are characterized by their ability to self-renew indefinitely and upon specific conditions differentiate into any cell type. These cells are maintained in specific growth medium to maintain their pluripotency and typically have a high proliferation capacity. We obtained two CDM1 mutant iPSC lines derived from CDM1 patient and one control line from a normal individual. We did not observe any obvious differences in their growth or appearance. For example, they all grew in compact colonies with round cellular morphology and with a high nuclear to cytoplasmic structure in bright field images (Fig. 1A). We further confirmed the presence of pluripotency/stemness markers including Sox2 and Oct4 by immunofluorescence and western blot (Fig. 1B, C) in all the three iPSC lines. Next, we performed a well-established southern blot hybridization method to determine the CTG triplet nucleotide repeat length in the three iPSC lines (Fig. 1D). While the repeat length/number quantification confirmed ∼1260 and ∼2020 CTG repeats for CDM1A and CDM1B cells respectively on the 3∼ end of the DMPK gene; there was no band/repeats observed in the CTL iPSC line (Fig. 1D). These results confirmed the presence of CTG triplet nucleotide mutation in the CDM1 lines and despite the mutation these lines displayed comparable growth pattern.

**Fig 1.**
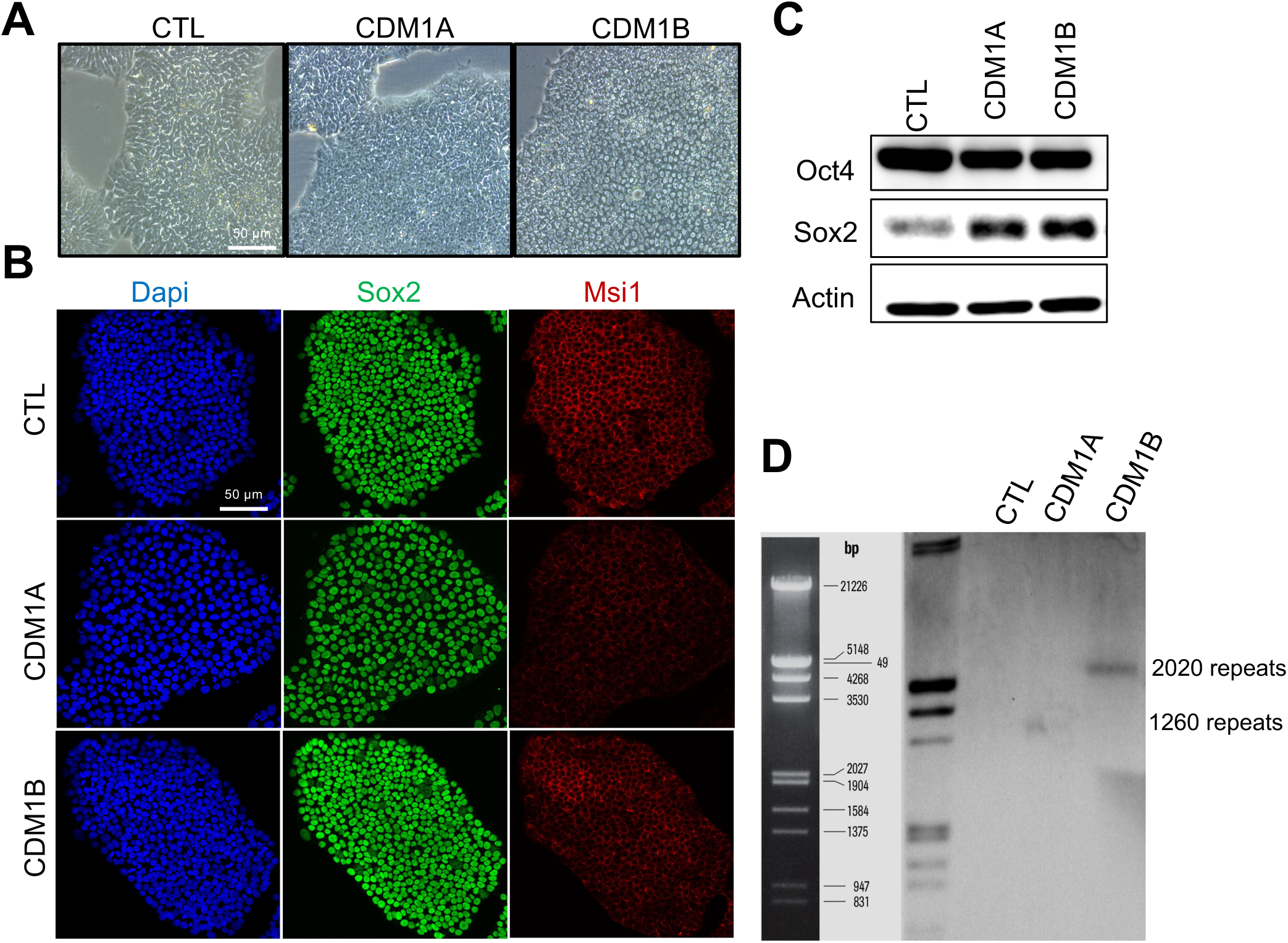
Characterization of control (CTL) and CDM1 (CDM1-A and CDM1-B) iPSC lines. (**A**) Bright-filed images of control and CDM1 iPSCs showing a similar pattern of growth, cell morphology cluster formation during maintenance. (**B**) Immuno-stained confocal images (20X) of control and CDM1 lines with DAPI and Sox 2 and Musashi1. (**C**): Western blot analysis for stem cell markers Sox2, and Oct4 during the maintenance of iPSC stage. Actin is used as the loading control. (**D**) Southern blot analysis showing comparative DMPK-CTG repeats in iPSCs. The blot shows bands corresponding to 1260 and 2020 CTG repeats for CDM1A CDM1B respectively while the control shows no indication of band intensity.

### NGN2-induced direct reprogramming revealed proneuronal differentiation by CDM1 mutants

Progressive cognitive impairment and neurodegeneration are the hallmarks of adult DM1 while CDM1 is categorized as a neurodevelopmental disorder (De Serres-Berard et al. 2021) (Sweere et al. 2023). To investigate any potential impact of *DMPK*-CTG^exp^mutations in neuronal differentiation and function, we used a well-established direct reprogramming approach to efficiently convert iPSCs into neurons (bypassing neural stem cell stage). In this approach, forced expression of neurogenin 2 (NGN2) and subsequent coculturing with mouse astrocytes efficiently generates mature functional neurons within 2-3 weeks (Fig. 2A)(Chen et al. 2020) (Zhang et al. 2013). Strikingly, within the first 7 days of differentiation, both CDM1 mutant lines displayed more mature neuronal morphology with elongated and elaborate processes/neurites compared to a modest neuron-like cells in the CTL (Fig. 2B). In addition, CDM1 mutants also generated more neurons compared to the CTL line as seen in bright field images (Fig. 2B). We then continued their differentiation and fixed and stained the cells at the end of 3^rd^ week (day 21) with various neuronal markers including Tuj1 (betaIII-tubulin) and Map2 (Fig. 2C). While both mutant lines showed a strong increase in Tuj1+ cells, overall Tuj1 intensity and Map2 intensity, only CDM1B showed no change in Map2+ cells compared to that of control line (Fig. 2D). Because TuJ1 expresses much earlier than Map2 during neuronal differentiation this data indicates a general stronger maturation phenotype in both lines with CDM1B being slightly delayed. To further explore the maturation stage of these cells, we stained them with two other mature neuronal markers including NeuN and synapsin (Fig. 2E-G). While NeuN is expressed by post mitotic neurons and is a pan-neuronal marker, synapsin is expressed later and is a marker of synapse forming neurons. Quantification of the synapsin and NeuN signal as % of Tuj1 positive cells among three iPSC lines showed no differences (Fig. 2G), indicating that the cells that have committed towards neuronal fate (TuJ1+) are all maturing to similar extent. This data also indicates that roughly 30% of TuJ1+ cells are maturing to form synapses (Fig. 2G). Taken together, these results suggest that mutants formed mature cells like control but with a higher number (Figure 2D).

**Fig 2.**
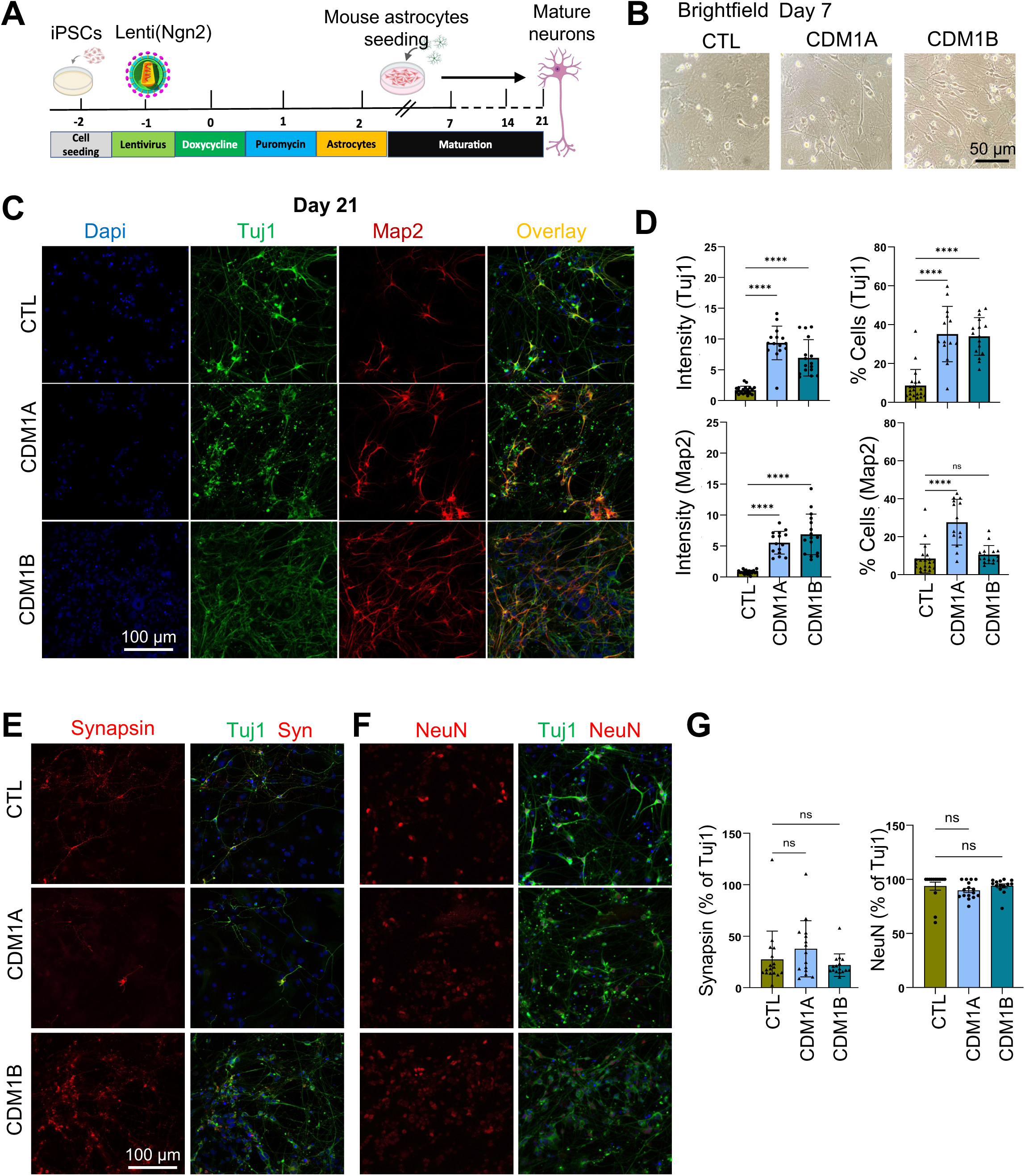
Direct reprogramming of CTL and CDM1 iPSCs to neurons using lentiviral mediated Ngn2 overexpression. (**A**) Schematics of the Ngn2 overexpression based direct reprogramming. (**B**) Representative bright filed images captured on the 3rd day during the selection process with doxycycline display long extensions of the processes and rosette formations in the CDM1 lines compared to the control. Representative confocal images (**E**) of Tuj1 (green) with Synapsin1 (red) or (**F**) of Tuj1 (green and NeuN (red) and their quantification (**G**) at 21 day in differentiation. Data represents ± SEM; n=15-20 images for (D); n=12-14 images for NeuN (G); n=15-17 images for Syn (G); from three independent cultures. n.s.= non-significant; **** = *p*<0.0001, one way ANOVA.

### CDM-1 iPSC lines form premature neurons during conventional neural induction

NGN2 direct reprogramming uses lentiviral delivery method which often integrates in the genome randomly, we suspected that the forced expression of exogenous Ngn2 together with some random integration might somehow alter cellular responses favoring rapid and mature neuronal formation in this method. Therefore, we decided to switch to conventional mode of inducing the iPSCs to neural progenitor cells and subsequently towards neurons. We used the commercially available neural induction medium supplemented with SMAD inhibitor to promote neural progenitor cells (NPCs) (Fig. 3A). First, we assessed the iPSCs conversion to NPCs at the end of the full neural induction protocol (3-week) using NPC marker nestin, Musashi 1 (Msi1) and Sox2 (Fig. 3B, C). We observed a very high efficiency of neural induction in this condition (Fig. 3B, C) however we also noticed nestin positive cells with elongated neurite like processes in CDM1 lines particularly so in CDM1A line (Fig. 3B). Because a small fraction of NPCs spontaneously differentiate into neurons and astrocytes in this neural induction protocol; we therefore stained these cells with neuronal (TuJ1) and astrocyte (GFAP) markers. Indeed, we observed a baseline spontaneous differentiation of Tuj1+ neurons and GFAP+ astrocytes in the CTL cultures (Fig. 3C). On the other hand, both mutant lines showed a high percentage of TuJ1+ cells with long neurites, while GFAP+ cells were absent (Fig. 3C). These data collectively suggest that CDM1 mutants are likely programmed towards neuronal fate at the expense of astrocytes. Next, we wanted to assess the timing of the precocious neuronal differentiation of CDM1 lines in this protocol. Therefore, we fixed and stained these cells at each passage (P1, P2 and P3) corresponding to ∼day 7, 14 and 21 in differentiation (Fig. 3D) with neuronal markers Tuj1 and Map2 (Fig. 3E,G). Analyzing these changes on a weekly basis, we observed that the spontaneous generation of Tuj1+ cells with neurite starts on P2 in CTL and does not further increase at P3 (Fig. 3E, F-top panel), while CDM1B showed appearance of these earlier at P1 and continued increasing through P3 and CDM1A showed appearance at P2 and kept increasing through P3 (Fig. 3E, F-top panel). Interestingly, quantification of average Tuj1+ neurite length further shows early emergence (at P1) and continued elaboration of neurites in CDM1 mutants compared to the CTL (Fig. 3E, F-middle panel). Once again, TuJ1 intensity quantification confirms the similar conclusion (Fig. 3F-lower panel). In line with this, Map2+ neurite length and Map2 intensity was also significantly higher in CDM1 mutants compared to CTL cells (Fig. 3G,H). Because the results from neural induction method (Fig. 3) were similar to that of NGN2-induced direct reprogramming method (Fig. 2) we conclude that mutant cells are programmed to form premature neurons and may be the underlying cause of CNS pathology in congenital DM1 patients.

**Fig 3.**
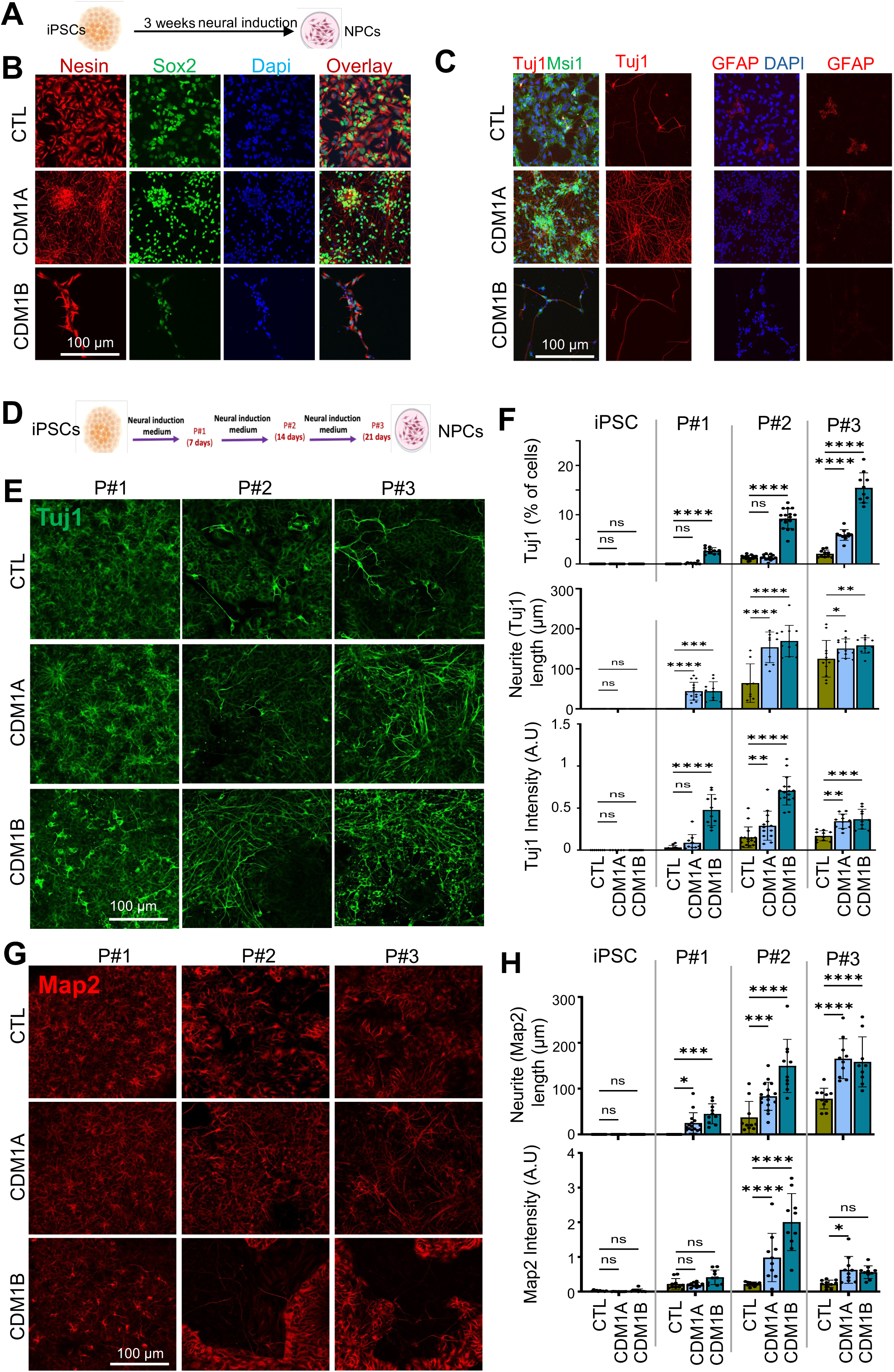
Neural induction reveals proneuronal differentiation capability of CDM1 cells. (**A**) Schematic representation of the timeline of neural induction. Representative confocal images of neural stem cell markers as assessed by Nestin (red) and Sox2 (green) (**B**) or neural lineage cell markers as assessed by Tuj1 (red), Mushashi 1(green) and GFAP (red) (**C**) at the end of 3^rd^ week, indicating an extensive neurite extensions and neuron differentiation et the expense of astrocytes GFAP+ cells in CDM1 lines. (**D**) Schematic representation of weekly investigation of neural induction process of iPSCs. Representative confocal images (**E**) and quantification (**F**) of Tuj1+ neural cells (green) with neurites. Representative confocal images (**G**) and quantification (**H**) of Map2+ neural cells (green) with dendrites. Data represents ± SEM from n=6-16 images per group (**F**, Tuj1), n=10-13 images per group (**H**, Map2), three independent cultures. n.s.= non-significant; * = *p*<0.05, ** = *p*<0.01, *** = *p* < 0.001, **** = *p*<0.0001, two-way ANOVA with Dunnett’s multiple comparison.

### The premature neuronal differentiation in CDM1 cells is accompanied by increased MBNL1 and MBNL2 nuclear expression and aggregation

Given the importance of MBNL1 and MBNL2 in DM1, we next examined the expression dynamics of these two proteins during neural induction of control and CDM1 cells. The western blot images disclosed a negligible to very low MBNL1 or MBNL2 expression at the iPSCs (Fig. 4A). Interestingly, both the proteins showed a gradual increase upon neural induction in all the 3 lines (Fig. 4A). Intriguingly, there was no difference in the MBNL1/2 levels between CTL and CMD1 mutants. Nevertheless, these observations indicate an important role of both MBNL1 and MBNL2 during the neuronal differentiation.

**Fig 4.**
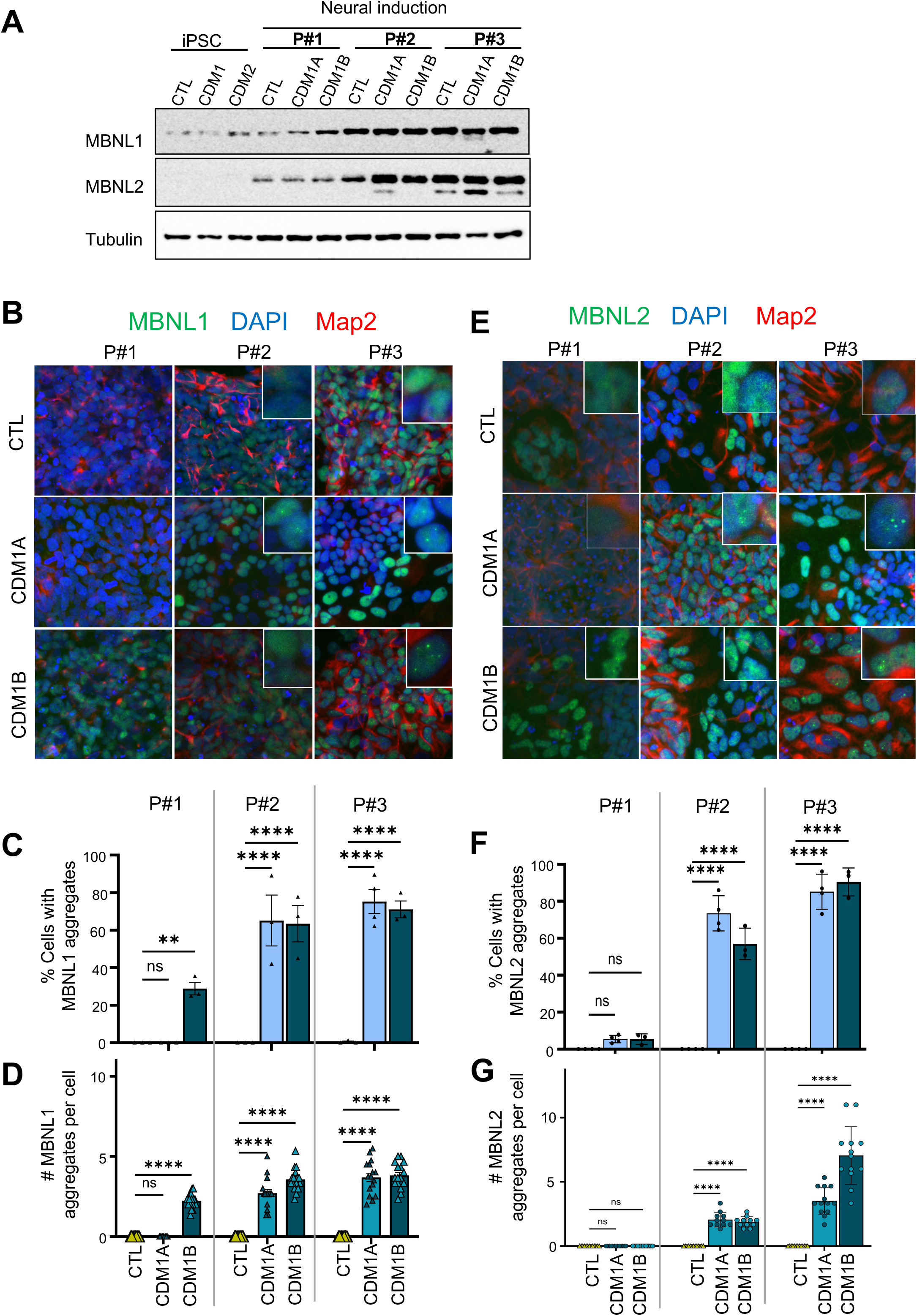
Expression analysis of MBNL1 and MBNL2 shows nuclear aggregation in CDM1 lines during neural induction. (**A**). Western blot analysis of MBNL1 and MBNL2 protein expression through 3 weeks of neural induction from iPSCs. Representative confocal images (63x) (**B**) and their quantification (**C,D**) shows robust nuclear aggregation of MBNL1 in CDM1 lines during the neural induction. Representative confocal images (63x) of MBNL2 (**E**) and their quantification (**F,G**) shows robust nuclear aggregation of MBNL2 starting at P2 during the neural induction. Images in the inset are enlarged view of aggregation of MBNLs. Data represents ± SEM from n=3-4 image per group from two independent cultures. n.s.= non-significant; ** = *p*<0.01, **** = *p*<0.0001, two-way ANOVA with Dunnett’s multiple comparison.

The hallmark of DM1 pathogenesis is the toxicity of the CUG repeats hypothesized to result from sequestration of RNA binding proteins that eventually disrupt their physiological function of RNA processing. MBNL1 and MBNL2 are also found to be aggregated in the brain neuronal cells of DM1 patients (Jiang et al. 2004) and motor neurons derived from DM1 iPSCs (Tahraoui-Bories et al. 2023). To examine aggregation (if any) of MBNL1 and MBNL2 in our cultures, we stained and performed high resolution confocal imaging (Fig. 4B, E). First, we found that MBNL1 and MBNL2 aggregates were present only in the CDM1 lines validating our CDM1 iPSC model. Second, more than 80% of all CDM1 mutant cells displayed aggregates in the range from 2 to 12 per nuclei while none detected in control cells (Fig. 4B-G). Moreover, the size and the intensity of the aggregates in the nucleus were intensified with the progression of neural differentiation. Interestingly, the earliest appearance of MBNL1 aggregates in CDM1B line at P1 (Fig. 4C,D) also matched with their earliest differentiation into TuJ1+ cell phenotype (Fig. 3F top panel). These results indicated a relationship between the loss of function of MBNL1 and/or MBNL2 and the mature neuron formation of CDM1 cells.

### Diverse neurogenic TFs, genes and mechanisms are accountable for premature neuronal phenotype in CDM1 lines

Since we observed the CDM1 mutant lines form neurons earlier and in abundance than the control cells, we desired to understand what factors and or mechanisms could propel this. For this, we performed RNA-sequencing studies using RNA isolated from the cells (control and CDM1 A&B) at the end of each passage. All libraries satisfied typical QC analysis and were sequenced to a depth to provide sufficient coverage analysis of gene expression and alternative splicing. Distinct transcriptomic profiles between the three cell lines were validated by the principal component analysis (PCA) and a correlation between the control and the CDM1 lines was made depending on the gene expression pattern (Figure 5 B). To observe any changes in the overall gene expression pattern, we combined P1, P2 and P3 sample RNA sequences for each cell line and compared with that of the combined iPSC RNA seq data, thereby providing enough sample size to perform stats for comparison between different groups. First, we identified the list of top 250 differentially expressed genes (DEGs) between neural-induced samples (total 9 samples; 3 each for three-iPSC lines) relative to iPSC (three) samples and performed the gene ontology for biological process (Fig. 5C). As expected, the GO showed neuron development/generation and brain development related processes (Figure 5C) confirming our successful neural induction. Next, we assessed the upregulated DEGs between the three groups during neural induction by the statistical significance of adjusted p-value (p ≤ 0.001) and subsequently used a 3-way Venn diagram to compare those (Fig. 5D). This way, we identified 29 DEGs shared between all three groups and 49 genes uniquely shared between CDM1A and CDM1B only (Fig. 5D). While gene ontology for biological process analyses identified several common processes including brain development and nervous system development in all three groups, glutamatergic neuron differentiation process appeared specifically in CDM1A and CDM1B shared genes suggesting mutant lines preferentially formed glutamatergic neurons (Fig. 5D). This diagram also showed that CTL and CDM1A cells shared the most no. of genes among them (Fig. 5D) corresponding to neuroblast polarity and membrane/dipeptide transport functions (Fig. 5D).

**Fig 5.**
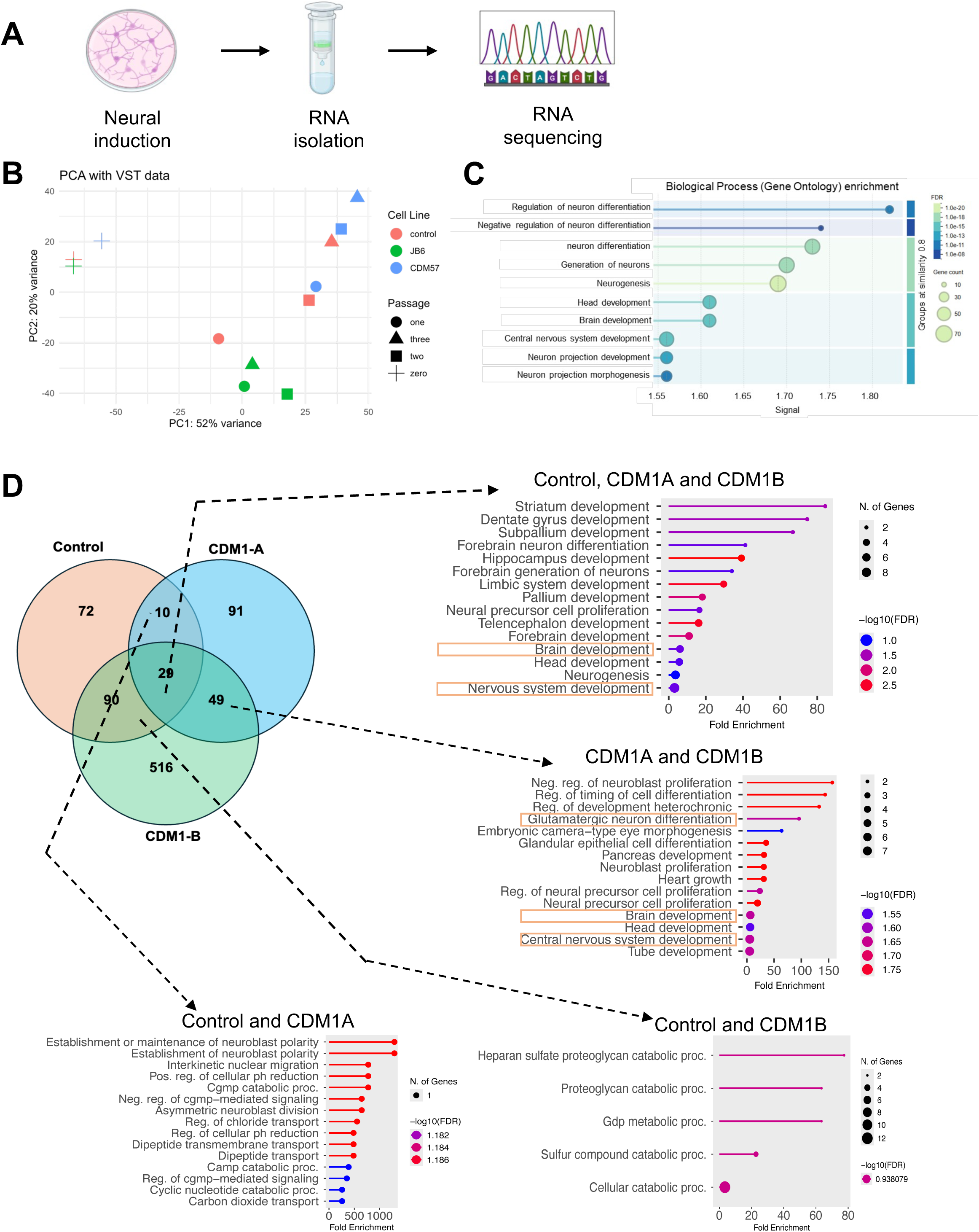
RNA seq analyses of CTL and CDM1 cells during neural induction. (**A**) Experimental workflow of RNA isolation, RNA sequencing during neural induction of control and CMD1 lines (total 12 samples). (**B**). Principal component analysis (PCA) plots of control and CDM1 lines based on the expression of genes during neural induction. The red, green and blue colors represent the control, CDM1A and CDM1B cell lines respectively. (C). Gene Ontology (GO) of biological process enrichment analysis of DEGs in the CDM1 and control cells after 21 days of neural induction relative to iPSC stage. (**D**). Venn diagram generated with upregulated genes in control and CDM1 lines as compared to their iPSC with an adjusted p value of ≤ 0.001. GO of biological processes enrichment analysis of shared genes between various groups using ShinyGO 0.85 gene ontology tool.

The early development of neurons in vertebrates is regulated by transcriptional factors (TFs) that control the proliferation, inhibition of pluripotency, expansion of neural blast cells and their transition to neural progenitors (Lee et al. 2014). Among these, basic-helix-loop-helix (bHLH) and other TFs are essential regulators of neural stem cell specification and differentiation during embryonic nervous system development (Dennis et al. 2019; Lee et al. 2022; Vainorius et al. 2023). Therefore, we manually analyzed our RNA-seq data for many select genes, including Ascl1, NeuroD1, NeuroG2, NeuroG1, Pax6, Notch2, and DLL3, that are well known to promote neurogenesis and neuronal differentiation (Fig. 6A). The normalized averaged FPKM values for these genes were plotted as heat map of fold change compared to the iPSCs (Fig. 6A). Interestingly, most of these genes are several folds upregulated in both CDM1 lines (Fig. 6A) compared to the control. Within this heat map we also plotted DMPK, MBNL1 and MBNL2 genes directly associated with DM1 pathology (Fig. 6A) which mostly remained similar between the groups. Using real-time PCR analysis, we analyzed the expression of the major bHLH family transcription factors including Ascl1, Neurog1, Neurog2, and NeuroD1 during neural induction. Interestingly, qPCR analyses confirmed that most of these neurogenic/neural differentiation transcription factors are upregulated in mix and match pattern in CDM1 lines validating our RNA-seq data (Fig. 5B-D). While upregulation of Ascl1, NeuroD1 and NeuroG1 was shared by both CDM1 lines (Fig. 6B), NeuroG1, Lmx1B and EBF3 generally was more induced in CDM1B line (Fig. 6C) and expression of PAX6, Lhx2 and FoxG1 was primarily increased in CDM1A line (Fig. 6D). Next our expression analyses of synapse associated genes including VgluT2 (Slc17A6) and GRIA2 identified an increase in both CDM1 lines (Fig. 6E) validating GO analyses from commonly induced genes (Fig. 5D). These results compelled us to further look at our RNA seq data for the expression of direct transcriptional targets of these transcription factors and are known to promote, neurogenesis, axonal growth, neuronal differentiation, migration, and development. Interestingly, we found these genes including, TBR1, NRCAM, Fezf2, NTRK2, NNAT, NR2F1, and NHLH1 to be upregulated in CDM1 lines (Figure 6A). These results indicated that the TFs which acts as a neurogenic switch and functions in controlled or regulated fashion are irregularly expressed in CDM1 lines that ultimately resulted in their early neuronal phenotype. Moreover, we also measured genes encoding neuronal cytoskeletal protein DCX and synaptic vesicle associated protein SNAP25(Fig. 6E,G).

**Fig 6.**
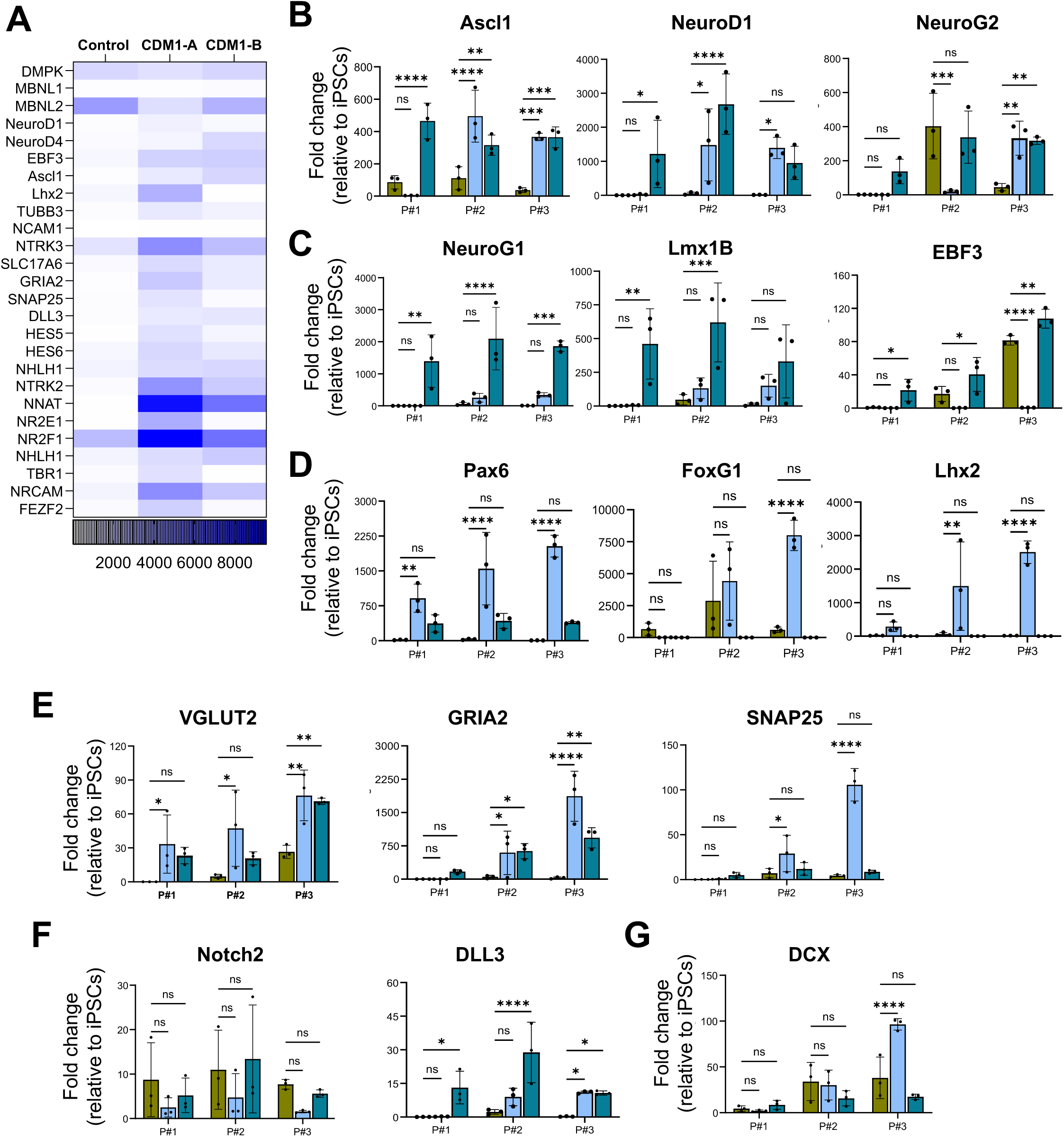
Upregulation of neurogenic genes and pathways in CDM1. (**A**) Heat map depicting average fold change in the expression of neurogenic genes obtained from CTL and CDM1 line RNA sequencing data during neuronal induction relative to iPSCs. (**B-G**) Quantitative real-time PCR analyses of select genes plotted as fold change in expression relative to their respective iPSC stage. Data represents ± SEM from n=3 independent cultures per group. n.s.= non-significant; * = *p*<0.05, ** = *p*<0.01, *** = *p* < 0.001, **** = *p*<0.0001

Because inhibition of Notch-signaling is one of the prerequisites for conversion of NPCs to early neuronal differentiation(Nelson et al. 2007; Kaltezioti et al. 2010), next we analyzed expression of Notch-signaling associated genes (Fig. 6F). Indeed, we found a trending downregulation of the Notch2 receptor expression and a strong and consistent upregulation of ligand DLL3, a notch-signaling inhibitor(Ladi et al. 2005), in the CDM1 mutant lines (Fig. 6F). Therefore, in addition to neurogenic transcription factor activity; Notch-signaling is also altered together leading to early neuronal differentiation phenotype in the CDM1 lines.

### MBNL1 and MBNL2 aggregation leads to changes in differential gene expression and alternative splicing activity in CDM1 lines

A leading cause of DM1 pathogenesis is mis-splicing of several genes due to aggregation and loss of function of MBNL group of proteins on expanded CTG repeat in DMPK mRNA. Next, we analyzed our RNA sequencing data from CDM1 and CTL lines to evaluate any changes in alternative splicing during their neural induction. Differential alternative splicing was analyzed using rMATS with a focus on skipped exon events based on junction count data with significant events defined by FDR<0.05, |ΔPSI|≥0.01, and >5-10 junctions reads in ≥70% of samples per group. The final ΔPSI was calculated as the average of ΔPSI of CDM1A and CDM1B relative to the CTL. These analyses identified a total of 392 splicing events. Out of total 392; 177 events were with a positive ΔPSI value and 215 events with a negative ΔPSI indicating increased exon inclusion and increased exon exclusion respectively.

Gene ontology (GO) analysis of biological processes of the differentially spliced genes revealed the GO terms involved in cell morphogenesis in neuron differentiation, neuron projection morphogenesis, generation of neurons, nervous system development, neuron differentiation and neurogenesis (Figure 7C) again confirming proneuronal phenotype of CDM1 lines. Further, the GO toward disease alliance analyses of these differentially spliced genes informed their association with Huntington’s disease (HD) (a rare inherited neurodegenerative disorder that affects the brain and nervous system) and autism.

**Fig 7.**
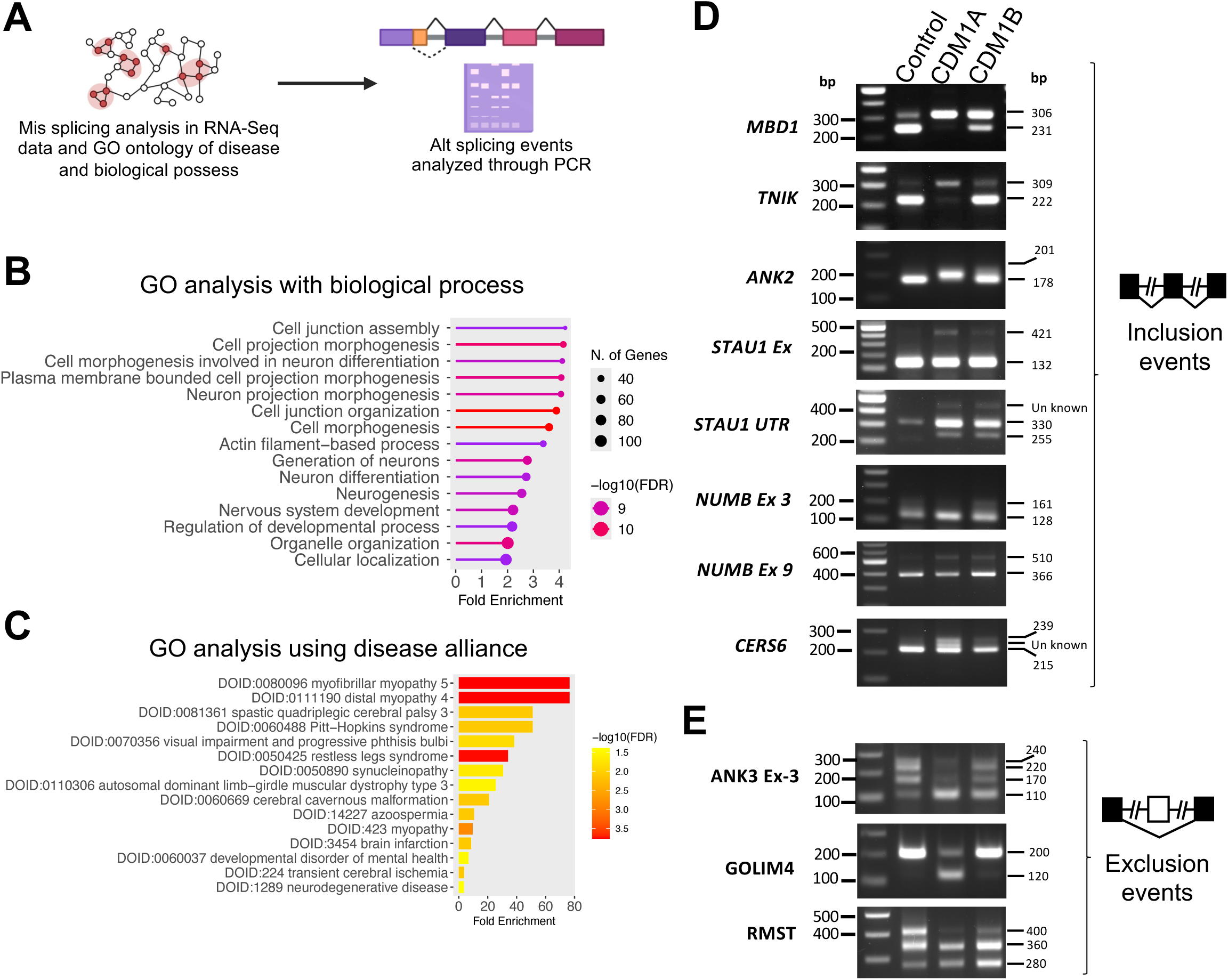
CDM1 lines display prominent mis-splicing during neural induction. (**A**) Schematics analyses of mis-splicing events from RNA seq data. (**B,C**) Gene Ontology analysis for biological process(**B**) and disease alliance (**C**) performed with a set of 392 mis-spliced genes identified using ShinyGO 0.85 version. (**D,E**) PCR analysis of selected mis-spliced genes with exon inclusion events (D) or exon exclusion events (E) in CTL and CDM1 lines at passage P3 of neural induction.

Since we observed a premature neuronal differentiation and aberrant expression of bHLH and other transcription factors in CDM1 lines, we sought to validate select number of genes for changes in alternative splice events by PCR analyses. We designed specific primers in such that PCR products would distinguish the inclusion or exclusion splicing events. The genes for splice analyses were selected based on their presence in the top 15-spliced event list and/or with known functions in neurogenesis and neuronal development including, MBD1, ANK2, TNIK, NUMB, GOLIM4, ANK3 and RMST. MBD1 (Methyl-CpG-binding domain 1) is a member of a family of methyl-CpG-binding proteins that function as epigenetic “readers,” connecting transcriptional control with DNA methylation (Zhao et al. 2003) and is known to regulate neurogenic switch in adult neural stem cells (Jobe et al. 2017). PCR results displayed a strong splicing disparity of MBD1 in CDM1 (Fig. 7B) indicating the critical role of MBD1 splice isoform in controlled neuronal differentiation. Similarly, splicing of ANK2 (encoding ankyrinB protein) is also greatly affected in CDM1 lines. Interestingly, mutations in ANK2 gene are associated with abnormal neurodevelopmental events leading to autism spectrum disorder, intellectual disability, and epilepsy (Yoon and Penzes 2025). Further, PCR analyses also confirmed splicing disparity in many other genes including, STAU1 (encoding an RNA binding protein); TNIK (encoding a serine/threonine kinase protein important in Wnt signaling regulation and neuronal synaptic plasticity) (Heraud-Farlow and Kiebler 2014) (Coba et al. 2012); NUMB (encoding a protein that antagonizes Notch signaling and controls neurogenesis and synaptic development) (Shao et al. 2017) (Sukhanova et al. 2025) and RMST (a brain specific long-non-coding RNA essential for neuronal differentiation) (Ali et al. 2024). These results suggest diverse mechanisms may be playing a role in CDM1 CNS pathology.

## Discussion

The development of the cerebral cortex is governed by precisely regulated, intricate processes that ensures accurate neuronal laminar positioning and the establishments of circuit-specific connectivity. Dysregulation at any step of this organized series of events, due to genetic mutations or environmental factors, leads to defined brain pathologies collectively known as malformations of cortical development (Guarnieri et al. 2018). Classical examples of such abnormalities include megalencephaly and hemimegalencephaly, conditions characterized by increased brain size in a symmetric or asymmetric manner and are now recognized as leading causes of drug-resistant epilepsy and intellectual disability (Guerrini and Dobyns 2014). Similarly, CDM1 infants display significant brain structural defects present at birth, consistent with CDM1 being a neurodevelopmental disorder (De Serres-Berard et al. 2021).

In this study, we discovered a link between the appearance of premature neuronal phenotype and the aberrant upregulation of neurogenic transcription factors in CDM1 cells. Although transcription factors ASCL1, NeuroG1, NeuroG2, NeuroD1 and EBF3 promote neurogenesis and neuronal development, they do so by acting sequentially at different stages. For example: ASCL1 is a pioneer transcription factor important in early neuroblast differentiation, NeuroG1 and NeuorG2 subsequently induce cell cycle exit and further neuronal specification while NeuroD1 promotes late neuronal development by inducing dendrite formation (Gao et al. 2009; Roybon et al. 2009; Aydin et al. 2019; Colasante et al. 2019; Bina et al. 2023). The marked expression of ASCL1, NeuroG1 and NeuroD1 in CDM1B cells at P1 (Fig. 6B,C) is consistent with neuronal differentiation and the emergence of dendrite-like structures, as corroborated by TuJ1 and Map2 immunofluorescence at P1 (Fig. 3E,G). Likewise, ASCL1 and NeuroD1 expression at P2 in CDM1A (Fig. 6B) coincides with the presence of neurons exhibiting neuritic processes (Fig. 3E-H). In contrast, the transcription factors Pax6, FoxG1 and Lhx2 modulate neurogenesis and neuronal development by acting in concert with NeuroG1, NeuroG2 and ASCL1, and by helping to maintain neural progenitor cells in a proliferative and undifferentiated state (Tian et al. 2012; Hou et al. 2013). The robust expression of Pax6, FoxG1 and Lhx2 in CDM1A throughout the entire neural induction phase indicates the presence of neural progenitor cells, which is further supported by the presence of rosette-like structures at P3 in CDM1A cells (Fig. 3B,C). Therefore, it is possible that CDM1 lines follow many different paths towards proneuronal phenotype as has been shown for the role of ASCL1 and NeuroG2 in converting mouse embryonic stem cells into neurons (Vainorius et al. 2023). Immunofluorescence based and or single cell RNA seq studies in future would help clarify this.

In this study we also found clues to possible signaling events that could promote proneuronal phenotype in CDM1 cells. Regulation of neurogenesis and neuronal differentiation through Notch pathway by Notch ligand Dll1/Dll3 is a well-recognized mechanism (Dennis et al. 2019; Lee et al. 2022). The increased expression of Dll3 in CDM1 lines (Fig. 6F) is in line with its known role in promoting neuronal differentiation by inhibiting Notch signaling (Ladi et al. 2005) an avenue that motivates future directions of inquiry in DM1. Interestingly, Dll3 is a direct target of Ascl1(Henke et al. 2009), which also were expressed at higher levels, in CDM1 lines (Fig. 6B). These results indicated the dysregulation of Notch signaling in CDM1 which is otherwise a guardian for controlled neurogenesis (Gonzalez and Reinberg 2025). Because same neural progenitor cells produce neurons first and later differentiate into astrocytes by a process termed as neurogenic to gliogenic switch in vivo (Miller and Gauthier 2007), and that CDM1 lines are proneurogenic (Fig. 2,3,4,5,6); it is imperative to ask whether this switch is affected in CDM1 lines? Interestingly, in our neural inductions, we almost never detected astrocytes (GFAP) in CDM1 lines while we always detected some spontaneous generation of astrocytes in the CTL line (Fig. 3C and data not shown). Therefore, we speculate that CDM1 mutations may affect this switch and promote neuronal differentiation at the expense of astrocyte differentiation. Indeed, a recent study showed impaired adhesion, spreading, and migration of astrocytes in mouse model of DM1 (Dinca et al. 2022).

The MBNL gene paralogs (*MBNL1*, *MBNL2* and *MBNL3*) are expressed in mice and humans. The spatial distribution of MBNLs within a cell varies in tissue and cell type specific manner (Kanadia et al. 2003; Sicot et al. 2017; Dinca et al. 2022). In line with this, while MBNL1 knockout mice showed a milder alternative splicing defect(Suenaga et al. 2012), MBNL2 knock out caused widespread alternative splicing changes in the brain (Charizanis et al. 2012). However, in our neural induction, both MBNL1 and MBNL2 expression levels increase similarly (Fig. 4A) and further both showed nuclear aggregation in CDM1 lines suggesting both may play roles in pathology. On the contrary, CDM1-B cells embraced MBNL1 nuclear aggregation much earlier than MBNL2, suggesting prominent role of MBNL1 during neurogenesis along with MBNL2. Although we did not include MBNL3 in our investigation, its overall contribution in DM1 pathology is neglected due to its low expression in adult muscle tissues. However, its role in different organs (Poulos et al. 2013; Choi et al. 2016), its expression in placenta, and its affinity to DMPK CTG mRNA repeats along with MBNL1 and MBNL2(Fardaei et al. 2002)suggest the possibility of higher percentage of maternally inherited CDM (Trucco et al. 2025) which needs to be explored.

In our study we detected aberrant splicing events of several genes. In addition, the gene ontology of biological and disease processes revealed their identity with cell morphogenesis and diseases relating to HD and autism related genes. This corroborates with a recent study where autism related traits in the mouse model of DM1 was due to MBNL sequestration and altered RNA splicing of autism-risk genes (Sznajder et al. 2025). Interestingly, while regulation of alternative splicing is one of the major functions of MBNLs, these proteins also impact mRNA localization, local protein translation and secretion of a subset of mRNAs implicating their varied roles in DM1 pathogenesis (PMID: 22901804). In line, we now provide evidence of dysregulation of neuron formation phenotype, nuclear aggregation of MBNL proteins, and the aberrant splicing events which may be collectively responsible for brain abnormalities in CDM1. Our data clearly reveals the cues for further studies and development strategies to combat CDM1 brain pathology.

## Conclusion

In conclusion, we demonstrate novel deficits in regulated neural induction of patient derived CDM1 iPSCs. These findings open new avenues for further mechanistic insights of CDM1 brain pathology and development defects. Our observation of continuous increase in neurons during each progressive week of neural induction is accompanied by increased intensity of MBNL1 and MBNL2 aggregation. Though we offer novel findings, there are few limitations that exist in our study. Our results are confined to few CDM1 patients derived and control iPSC lines and need to be extended to many patient derived iPSC lines to support our findings. Although both CDM1A and CDM1B cells generally showed a proneuronal phenotype, their differentiation dynamics was different (Fig. 6C-E) and could be attributed to patient specific inherent variations or due to their differential CTG repeat lengths (Fig. 1D). Nevertheless, our findings strongly implicate a model towards personalized medicine where patient specific treatment is devised based on patient specific disease mechanism/phenotypes (Hampel et al. 2023). Moreover, the robustness of our observations would be strengthened if these gene alterations are similarly observed in histological brain tissues derived from CDM1 patients. Extending our studies to isogenic lines by correcting the CTG repeats using novel gene editing tools would stipulate confidence in treating CDM1 conditions. Finally, understanding the fundamentals of DMPK mutations and additional reasons for CDM1 pathology would provide a large picture of the disease and open possibilities for treating this lethal disorder at embryonic stages that can preserve lives.

## Authors’ contributions

S.C.R.T. planned and performed most experiments, with assistance from S.M, J.P.G., C.T., and O.D. S.K.S. acquired funding. S.K.S. conceived the study, contributed to planning of the experiments and performed experiments. S.C.R.T. and S.K.S. drafted the manuscript. All authors read and approved the final manuscript.

## Acknowledgements

Special thanks to Nicholas Johnson, Melissa Hale, Marina Provenzano, Dove Enicks (Department of Neurology, Virginia commonwealth University, Richmond, Virginia, USA) for providing iPSC lines and assistance in RNA sequencing. We also thank Julia Hartman and Samuel Carrel (Department of Neurology, Virginia commonwealth University, Richmond, Virginia, USA) for their assistance in southern blot experiments.

## Availability of data and material

The data that support the findings of this study are available from the corresponding author upon reasonable request.

## Ethics approval

Mice were housed at Virginia Commonwealth University according to guidelines of the Institutional Animal Care Use Committee (IACUC). The mouse protocols were approved by the IACUC. All mice were housed with food and water available ad libitum under a 12 h–12 h light–dark cycle in a 20–22°C and 40–60% humidity environment.

